# Grow now, pay later: when should a bacterium go into debt?

**DOI:** 10.1101/2023.07.26.550682

**Authors:** Jaime G. Lopez, Amir Erez

## Abstract

Microbes grow in a wide variety of environments and must balance growth and stress resistance. Despite the prevalence of such trade-offs, understanding of their role in non-steady environments is limited. In this study, we introduce a mathematical model of “growth debt”, where microbes grow rapidly initially, paying later with slower growth or heightened mortality. We first compare our model to a classical chemostat experiment, validating our proposed dynamics and quantifying *Escherichia coli*’s stress resistance dynamics. Extending the chemostat theory to include serial-dilution cultures, we derive phase diagrams for the persistence of “debtor” microbes. We find that debtors cannot coexist with non-debtors if “payment” is increased mortality but can coexist if it lowers enzyme affinity. Surprisingly, weak noise considerably extends the persistence of resistance elements, pertinent for antibiotic resistance management. Our microbial debt theory, broadly applicable across many environments, bridges the gap between chemostat and serial dilution systems.

**Teaser:** Microbes can sacrifice future growth for immediate gains, how does this trade-off shape the structure of microbial communities?

## Introduction

Microorganisms have shaped the world we live in and have adapted to thrive almost anywhere, including the human body, desert soils, forest soils, and coral reefs (*1–4*). Competition for limited resources is a central characteristic of microbial life, and this fierce competition, ongoing since the first cell emerged billions of years ago, has led microbes to explore astonishing ways to gain a growth advantage. However, our understanding of the core principles behind microbial competition remains mostly limited to steady environments.

In the microbial world, organisms commonly make a trade-off between hedging against future adverse events and maximizing immediate growth (*5*). This trade-off manifests in a variety of ecological contexts, many of them relevant to human health. For example, carrying the genes necessary to resist antibiotics slows a microorganism’s growth in the absence of antibiotic, but the organism’s growth will be maintained if antibiotics enter the environment (*6–9*). Similarly, the malaria parasite *Plasmodium falciparum* incurs a substantial growth defect by carrying a mutation that protects from anti-malarial drugs (*10*). This trade-off is not limited to drug resistance: mutations in regulatory elements, such as *Escherichia coli*’s *rpoS*, allow organisms to trade-off nutritional competence and general stress resistance (*11, 12*). To understand the core principle behind this “grow now, pay later” phenomenon, we introduce the metaphor of microbial debt—in which lacking the response capacity to future adverse events, reaps an immediate speedup in growth that is then “paid back” in the future. So, debtors are species that shift cost to the future (similar to interest paid on a loan) to gain a benefit in the present. Yet, despite the ubiquity of debt-like mechanisms in nature, theoretical understanding of debt in non-steady environments is missing.

Debt causes a shift of cost from the present to the future, and its effect is thereby strongly tied to temporal variation. Temporal variation is a known driver of eco-evolution, impacting systems ranging from the microbiota of the Hadza people of East Africa (*13*), the mucosa of mice (*14*), and antibiotic resistance evolution (*15*). Evolutionarily, temporal variation can reduce the efficiency of natural selection and increase the fixation probability of mutations (*16*). Perhaps the simplest, yet most experimentally prevalent, example of temporal variation is serial-dilution batch culture - in which microbes are subject to periodic dilution and nutrient replenishment. This is substantially different from the chemostats — another common experimental system in which nutrient supply and dilution occur continuously (*17*). Studying and comparing the behaviors of chemostat and serial dilution systems, which represent two extremes of environmental variation, can provide insights into the environmental fluctuations present in real ecosystems. Although serial-dilution culturing is ubiquitous in experimental work (*18, 19*), modeling efforts in this direction are still incomplete (*20–23*). In the context of serial dilution cultures, we have recently shown that the effect of temporal variation can be understood through the “early-bird” effect (*20*). The early-bird effect arises from the temporal gap between the introduction of nutrients and the inoculating mixture of microbes (at *t* = 0, simultaneously); and the transfer of a fraction of the culture to inoculate the next batch (a dilution event, at *t* = *t*_*f*_). During this time, the early-bird species exhibit rapid growth at the onset of the batch cycle. Although their per-capita growth diminishes later in the batch cycle, the significant size of the ‘early-bird’ species can still consume nutrients, consequently inhibiting the growth of ‘late-bird’ species. This effect can cause shifts in community structure, shifts that do not occur in an equivalent chemostat. The early-bird effect has recently been shown to control the assembly of gut microbiome-derived *ex vivo* communities (*24*). It suggests a possible advantage, unique to serial-dilution cultures: to pursue a ‘debt’ or ‘grow now, pay later’ strategy which accelerates early growth by shifting costs to the future.

Much theory has focused on the dynamics of debt trade-offs in fixed, chemostat-like conditions, showing that that debt trade-offs can enable coexistence of strains (*25, 26*). In some cases, predictions from these chemostat theories have been qualitatively verified experimentally (*27*). While these trade-offs have been explored extensively in steady environments, less is understood about their dynamics in fluctuating environments. Contemporary understanding of the ecology of antibiotic resistance suggests that variation can substantially alter the value of debt (*28*), highlighting the need for further work. Biophysical theories have been developed that predict optimal growth behaviors of microbes under environmental fluctuations, frameworks which can be applied to stress resistance phenotypes (*29, 30*). However, despite recent progress (*31*), there remains a pressing need for a comprehensive theoretical framework for trade-offs in serial dilution systems, bridging across commonly used experimental setups.

In this manuscript we ask: what are the consequences of accruing debt for a microbe within the context of a consumer-resource model? How does debt impact the growth dynamics of microbial communities under serial-dilution conditions compared to those in chemostat environments? We develop a modeling framework to explore microbial debt and demonstrate its use to analyze previously published experimental data. Our theory makes no assumptions about the shape of the debt trade-off surface (*9, 27, 32*). Though we consider only fixed and unchanging trade-offs, it is possible to extend the model to account for non-genetic adaptation (*33*). Our proposed model mimics a typical serial-dilution experimental procedure, while also being directly applicable to chemostats. We study the competition between two species, a debtor and a non-debtor, and find that when debt is compensated for by increased mortality, coexistence cannot occur between debtor and non-debtor, regardless of whether the mortality occurs in a punctuated or continuous manner. However, when debt is paid through decreased metabolic enzyme affinity, a stable coexistence state emerges. Intriguingly, when the cost of debt is subject to noise, the losing species can survive for a much longer time when compared to an equivalent deterministic system. Therefore, noise can enhance the persistence of antibiotic resistance elements.

## Results

### Chemostat dynamics reveal growth-debt regulation in *E. coli*

Before introducing our serial dilution model, we first present the equivalent chemostat model as a baseline for the serial dilution results and as a bridge to the existing, chemostat-dominated literature. A single nutrient is provided at flux *S*, and its concentration within the chemostat is denoted *c*(*t*). The chemostat is well-mixed and constantly diluted with dilution rate *δ*. Let Species N be a non-debtor (a microbe with resistance to the stressor) with biomass density *ρ*_*N*_ (*t*). Nutrient uptake is controlled by *g*[*c*(*t*)], and growth is proportional to nutrient uptake. The per-capita growth rate of Species N is *E g*[*c*(*t*)] where *E* defines the maximal per-capita growth rate. In contrast, Species D, the debtor species, with biomass density *ρ*_*D*_(*t*), has higher maximal growth rate, *E* + Δ*E* due to sacrificing its stress resistance capability and diverting these resources to immediate growth. The lack of stress resistance leads to constant stress-induced death at rate *ω*. The equations governing these chemostat dynamics are therefore: 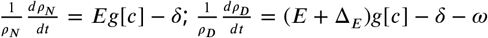; and 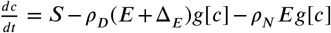. Note that we neglect *δc* in the nutrient dynamics as the dominant nutrient depletion term comes from the bacterial consumption (*25, 34*).

To validate our model assumptions and quantitatively show the existence of debt trade-offs, we apply this chemostat model to the classical experiments of Notley-McRobb et al. (*11*). These experiments studied competition between wild-type (WT) *Escherichia coli* and mutants with defective *rpoS* genes, the transcription factor governing stress response in *E. coli*. Here, the WT plays the role of non-debtor and the mutants are debtors. In ***Figure 1***A, we show the relative abundance of the non-debtor WT strain in chemostats operating at different dilution rates. In these low-stress chemostats, the community is rapidly overtaken by the debtor *rpoS* mutants, well-captured by our model fit (see SI Appendix ***section 4*** for details). However, these debtors have traded growth rate against stress resistance, and are now far more vulnerable to stress. In ***Figure 1***B we show the temperature stress response of the community from one chemostat as the debtor population increases. Initially, the population is composed of non-debtors and many bacteria survive the stressor. However, once debtors sweep the population almost no bacteria survive the stressor. These stress survival dynamics are well-described by our model fit. Thus, debt trade-offs occur within *E. coli* and are quantitatively consistent with our model assumptions

**Figure 1.**
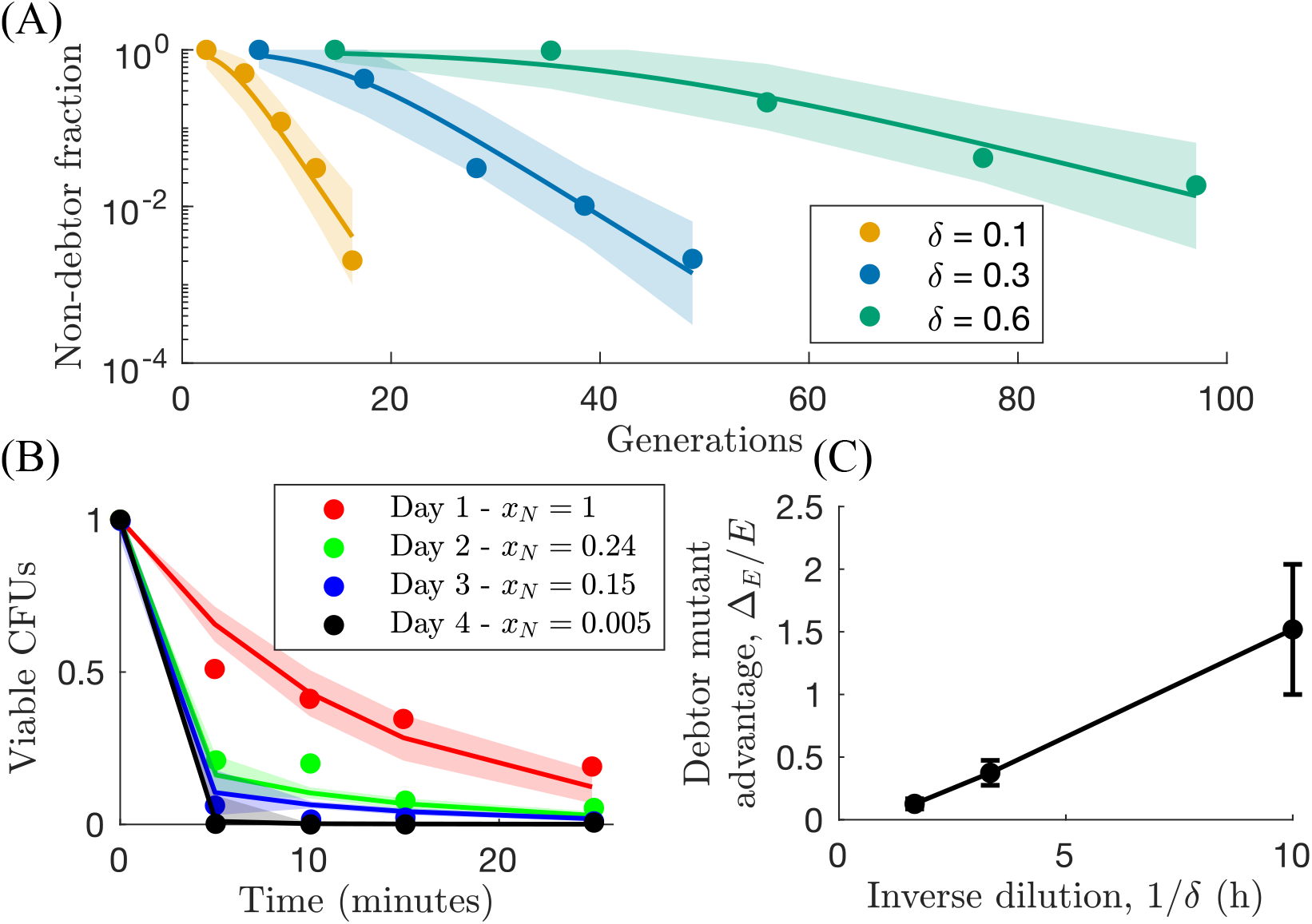
Application of chemostat theory to experiments of Notley-McRobb *et al*. validates model assumptions and shows existence of trade-offs between maximal growth rate and stress resistance in *E. coli* (*11*). (*A*) Fit of chemostat theory to *rpoS* evolution trajectories in pH 7 glucose-limited chemostats at three different dilution rates *δ*. Fit parameters: Δ_*E*_ /*E* = [1.52 ± 0.52, 0.37 ± 0.1, 0.13 ± 0.04] at *δ* = [0.1, 0.3, 0.6] h^−1^, respectively and *x*_*D*_(*t* = 0) = 0.025. (*B*) Fit of theory to community temperature stress assay at 60°C. Samples taken from glucose-limited *δ* = 0.3 h^−1^ chemostats at different days. Fit parameters: *ω*_*N*_ = 0.084 ± 0.014 min^−1^ and *ω*_*D*_ = 0.99 ± 1.92 min^−1^. (*C*) Plot of debtor mutant advantage Δ_*E*_ /*E* as a function of inverse dilution rate 1/*δ*. Values from fit in *A*.

In this chemostat formalism, when is growth debt worthwhile? For this calculation, we define *g*[*c*] as the classical Monod function 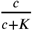, where *K* is the half-saturation coefficient. By comparing the competitive ability of the non-debtor and debtor, we determine that a stressor death penalty *ω* is worthwhile if the corresponding benefit Δ_*E*_ satisfies:

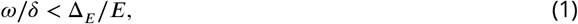

(for details see SI Appendix ***section 1***). Thus, at a higher dilution rate, *δ*, the relative cost of debt decreases. ***Equation 1*** predicts a linear relationship between inverse dilution rate 1/*δ* and the growth advantage Δ_*E*_/*E*, consistent with the debtor mutants observed by Notley-McRobb (*11*). In ***Figure 1***C, we show the growth advantage of debtor mutants as a function of inverse dilution rate. The debtor mutant Δ_*E*_/*E* is a reflection of *E. coli*’s investment in stress resistance. A large debtor advantage indicates that the non-debtor invests more resources in stress response, while a small advantage indicates little investment. The linear relationship shown in ***Figure 1***C indicates *E. coli* invests less in stress resistance at higher growth rates in a manner consistent with our theory.

Similar chemostat theories to those presented here have been further developed. For example, it has been shown that with additional information about the form of the *rpoS* trade-off surface, one can qualitatively predict the outcome of *E. coli* evolution within a chemostat (*27*). However, what happens away from chemostat conditions in a serial-dilution, fluctuating environment?

### Serial dilution model formulation

We now establish our serial dilution model where two species compete for a single, limited nutrient in an environment influenced by a stressor. In a serial dilution culture, microbes are grown in a well-mixed environment for a given period of time, *t*_*f*_, before being diluted into fresh media and allowed to grow again. We refer to the new nutrient added after dilution as the ‘nutrient bolus’. A single type of nutrient is provided, and its concentration within a batch is denoted *c*(*t*) with *c*(*t* = 0) = *c*_0_. As in the chemostat, let Species N be a non-debtor (a microbe with resistance to the stressor) with biomass density *ρ*_*N*_ (*t*), with per-capita growth rate *E g*[*c*]. In contrast, Species D, the debtor species, with biomass density *ρ*_*D*_(*t*), has higher maximal growth rate, *E* + Δ*E* due to sacrificing its stress resistance capability. The lack of stress resistance leads to two possible forms of stress-induced death: a continuous death rate per unit time (*ω*), and/or, an increased dilution during dilution events (Ω). The within-batch dynamics are therefore:

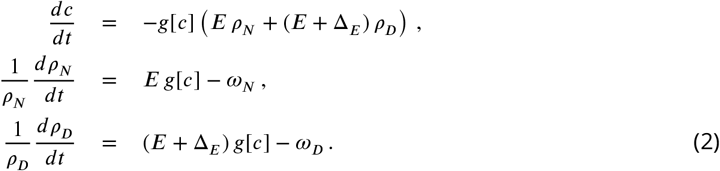

Here, *ω*_*N*_ and *ω*_*D*_ are the non-debtor and debtor within-batch death rates, respectively. These death rates could be the result of an antibiotic in the media or another stressor, such as temperature, salinity, or pH. Note that the terms on the right hand side of the equations for *ρ*_*N*/*D*_ are the percapita growth rates. It is possible to render these equations dimensionless by measuring time in units of *E*, and concentration in units of *K* (*cf*. SI Appendix ***section 3***). However, since our model is easily applicable to experimental data, we keep the dimensionful version of these dynamics for more direct data comparison.

The second source of mortality in the model is not continuous, but rather, punctuated, happening only at the transition between batches. We refer to the transition from batch to batch as ‘dilution’, reflecting what occurs in laboratory serial dilution, but it represents any number of potential punctuated stressors, such as sudden heat or osmotic shock. When dilution happens, each species is diluted by its species-specific factor Ω_*N*_ or Ω_*D*_. Thereby, dilution in our model generalizes the typical scenario where a small volume of a well-mixed culture is used to inoculate the next batch, diluting each species equally. In this generalized version, the total inoculum size, *ρ*_*N*_ (0) +*ρ*_*D*_(0), can vary from batch to batch until steady state is reached. A schematic of these mechanisms is shown in ***Figure 2***A. At the end time (*t*_*f*_) of batch *b*, each species gets diluted differently, and so, at the beginning of batch *b* + 1,

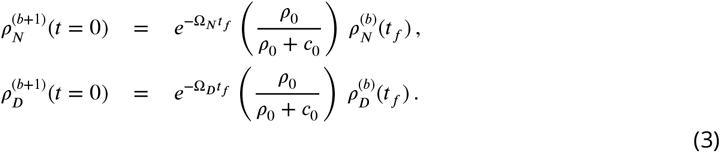

In the rest of this manuscript, we set Ω_*N*_ = *ω*_*N*_ = 0 and drop the subscript from Ω_*D*_ and *ω*_*D*_.

**Figure 2.**
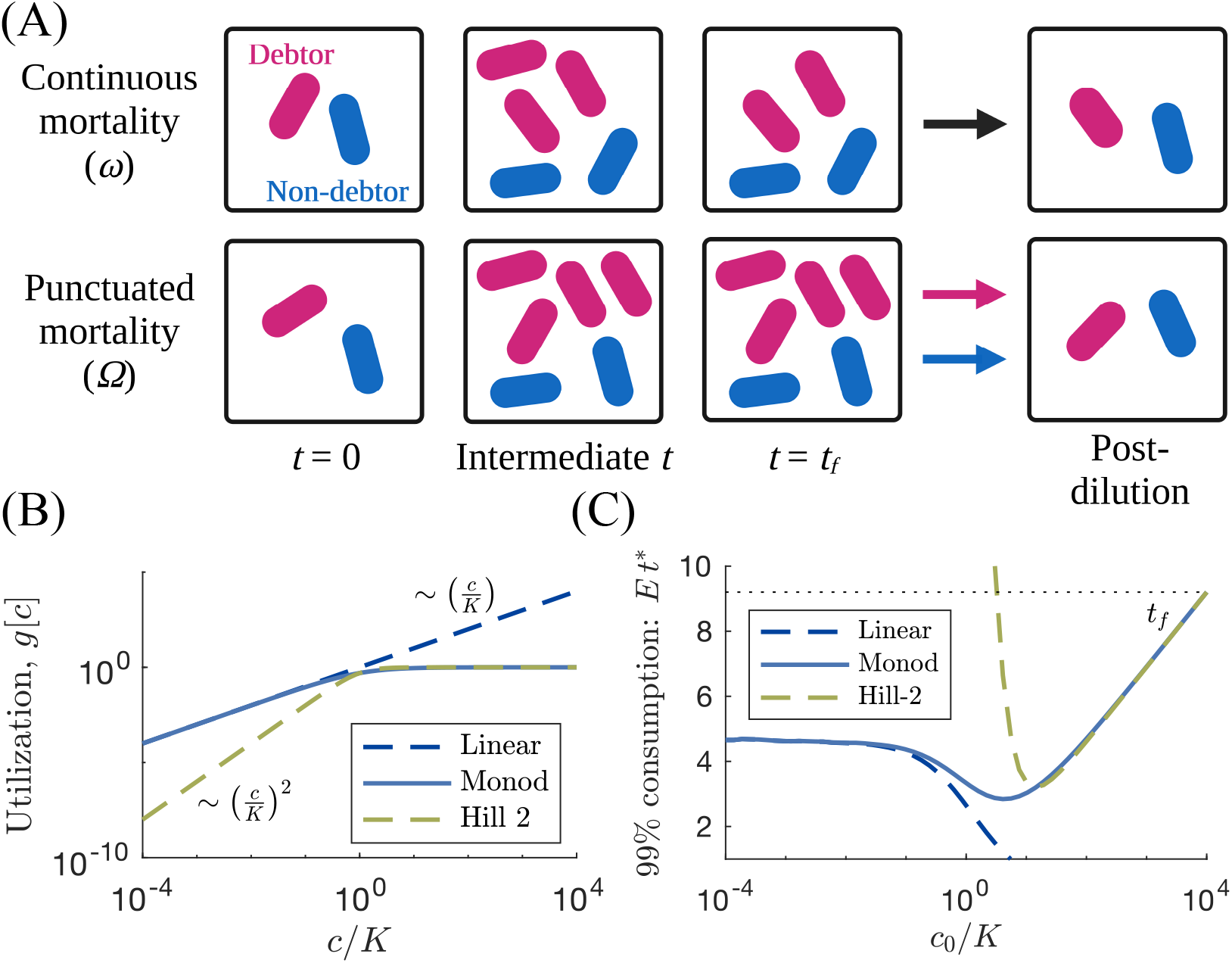
Overview of serial dilution ecosystem with debt. (*A*) Schematic of the serial dilution system with debt. In the continuous death case, the debtor is able to initially grow more quickly, but loses biomass continuously. In the punctuated death case, the debtor grows more quickly but is diluted more severely at the end of the batch. (*B*) Example of growth functions. Linear is *g*[*c*] = *Ec*, Monod is 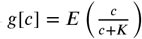, and Hill-2 is 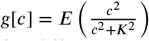. (*C*) Time necessary for a single non-debtor species to deplete 99% of the supplied nutrient *t*^∗^for different growth functions. In order to choose a single *t*_*f*_ that is sufficiently long, we select the *t*^∗^ for *c*_0_/*K* = 10^4^ with a Monod growth function.

#### Nutrient utilization, g[c(t)]

We considered two general forms of the utilization function *g*[*c*]: utilization that is proportional to the nutrient concentration, *c*(*t*), and utilization that obeys Hill-like non-linearity. Specifically,

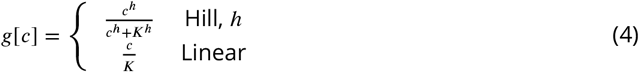

The *h* = 1 case is known as the Monod function (*17*), and its *c* ≪ *K* limit corresponds to linear consumption. At the opposite limit, at high nutrient concentration, linear consumption is nonphysical as it implies that a huge quantity of nutrient would be consumed almost instantly. Additionally, if *c* ≪ *K* and *h* > 1, at low nutrient levels, nutrient consumption effectively grinds to a halt. The Monod case, *h* = 1, extends between these two behaviors and is well-established to model microbial nutrient utilization (***Figure 2***B). For the rest of this manuscript, we thus focus on the Monod form.

#### Batch time, t_f_

When modelling serial dilution, it is possible to choose the batch time *t*_*f*_ to be as long as necessary to consume all resources (*20, 21*). However, here we also considered within-batch mortality, and therefore arbitrarily long batches would lead to the extinction of the susceptible species. A more realistic view, in line with experimental practices, sets a fixed batch time, *t*_*f*_. The time needed for consumption of most of the nutrient depends on how much nutrient is provided. In this study, we varied the supplied nutrient concentration, *c*_0_, over six orders of magnitude. To ensure that most of the nutrient is consumed in all relevant scenarios, we define *t*_*f*_ based on the nutrient utilization at high *c*_0_. Assuming Monod utilization with only one species present, *t*_*f*_ is defined as the time it takes to consume 99% of *c*_0_ with *c*_0_/*K* = 10^4^. Thereby, we ensure that at least 99% of the nutrient will be consumed for all relevant scenarios explored in this work. The dependence of *t*_*f*_ on *c*_0_, while maintaining 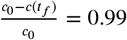, is shown in ***Figure 2***C.

### Equivalent value of debt represented as continuous and punctuated mortality

In our model, when is microbial debt worthwhile? Does it matter if the debt incurred is paid for in a continuous or punctuated fashion? We quantify debt by considering the dynamics within a single batch, i.e., a single growth and dilution cycle. At a given batch *b*, it is possible to integrate Eq. 2, given the initial inocula concentrations 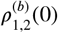. At the end of this batch, *t* = *t*_*f*_, we have the following populations:

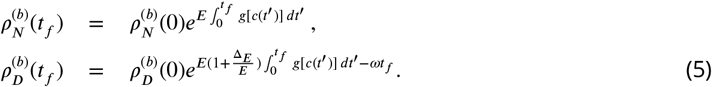

When going to the next batch, the debtor species gets diluted by a multiplicative factor 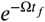 beyond the 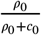 factor both species get diluted by. Therefore, at the beginning of batch, *b* + 1

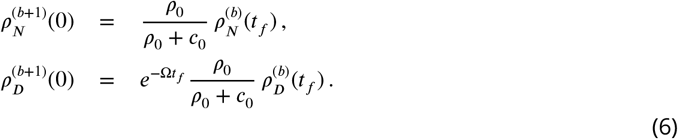

At steady state, equal growth implies that both species are increased and decreased by the same fold-change. Therefore, coexistence implies that,

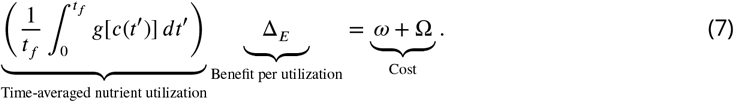

Equation 7 states that coexistence balances growth cost with benefit: the growth benefit per nutrient utilized times the time-averaged nutrient utilization equals the cost. In essence, the benefit from accruing the debt in the form of increased growth must be equal to the death cost incurred. This expression defines the border between regimes in which debt is favorable and those in which it is unfavorable (***Figure 3***A). In this choice of parameterization, the two forms of debt, continuous (*ω*) and punctuated (Ω), are equivalent and their rates are additive.

**Figure 3.**
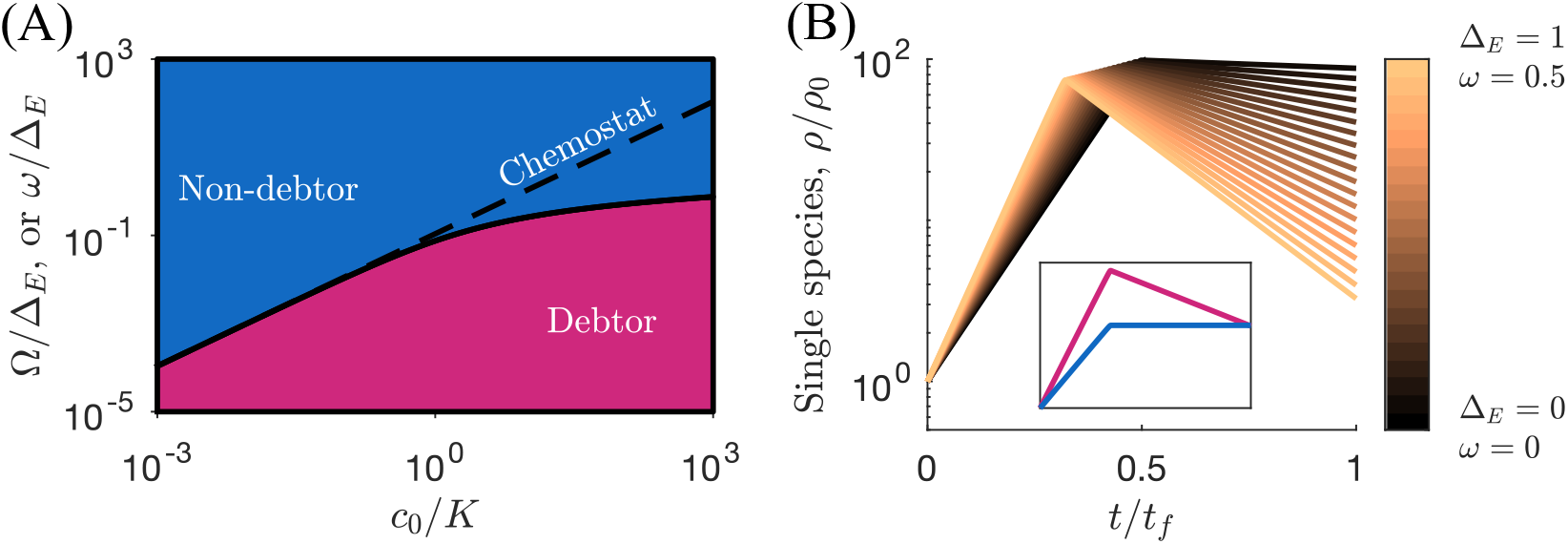
Serial dilution results in (*A*) Phase diagram of competition between debtor and non-debtor as a function of bolus size *c*_0_/*K* and normalized death penalty. The dashed line represents the phase boundary for the equivalent chemostat system, diverging substantially from the serial dilution phase boundary at high *c*_0_/*K*. This phase diagram captures both continuous and punctuated death, with the *y*-axis being equivalently Ω/Δ_*E*_ or *ω*/Δ_*E*_ (*B*) Within-batch dynamics of a single species along the Δ*E, ω* trade-off. Note that while the debtor species grows rapidly initially, its final biomass may be much lower due to death. *Inset*: (same axes). In competition with a non-debtor, the increased early growth of the debtor consumes the nutrient and denies nutrient from the non-debtor.

Importantly, in the single species case, even away from steady state, the nutrient utilization integral is independent of *g*[*c*] if the vast majority of the nutrients are consumed within the batch, with 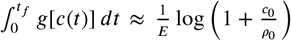 by mass balance. Note that 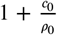 is the dilution factor of the serial dilution system, a property determined by the experimentalist. This is a tremendous simplification, since knowledge of *g*[*c*] is unnecessary. It is especially useful to parameterize experimental mono-culture dynamics, without having to measure *g*[*c*]. For example, application to an antibiotic resistance measurement requires only the element’s carriage cost and the death rate of cells without the resistance elements.

### Serial dilution cultures show diminishing return on debt at high nutrient concentrations when compared to a chemostat

We have determined the phase diagram for debt favorability in a serial dilution system, but how does this compare to a system with continuously supplied nutrient, i.e., a chemostat? If we add a very small bolus, *c*_0_ ≪ *K*, then dilute the culture by a fraction close to 1, and repeat, it stands to reason that the limiting behavior becomes continuous in time. As *c*_0_/*K* → 0, the serial dilutions become effectively a chemostat, and so, we expect that the continuous and punctuated dilution cases have the same chemostat limit. To consider *c*_0_ ≪ *K*, the chemostat limit, we follow (*20*), and linearize the dynamics around the small parameter *c*_0_/*K* (for details see the *SI Appendix* ***section 4***). The chemostat cost-benefit tradeoff then becomes 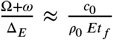, shown as the dashed line in ***Figure 3***A. It is satisfying to note that this is equivalent to the linear chemostat debt relation in Equation 1 and thus also consistent with the experimental analysis shown in ***Figure 1***C.

Why does the boundary between debtor and non-debtor dominance in the serial-dilution culture deviate from its equivalent chemostat at large *c*_0_? There is a diminishing return on debt as more nutrient is supplied. The diminishing return and its relation to the early-bird effect can be understood from ***Figure 3***B, showing time-series of a debtor in a single batch, grown in mono-culture, with several values of (*ω*, Δ_*E*_) pairs. The pairs lie along the coexistence boundary with the non-debtor (***Figure 3***A). As *ω*, Δ_*E*_ are increased, the species grow more rapidly initially, (higher slope in the semi-log plot), then followed by an exponential loss because of *ω* (downward slope). The initial debt, which gives the rapid rise, is offset by the incurred death later in the batch. Indeed, when the non-debtor species is introduced in a co-culture, both species reach the same level at the end of the batch (inset of ***Figure 3***B). This demonstrates the early-bird effect since the debtor deprives the non-debtor from nutrient in the early growth phase, though finally the debtor settles at a much lower population size because of the debt payoff.

### Debts involving a growth affinity trade-off allow coexistence

Thus far, we have examined trade-offs where the maximum growth rate under normal conditions is counterbalanced by mortality in the presence of a microbicidal stressor. However, there are other forms of growth stress and payment, e.g., there exist microbiostatic stressors that do not kill but rather inhibit the growth of microbes. The inhibition can be considered as a penalty on the maximum growth rate. Similarly, resistance mechanisms can affect growth properties other than the maximum growth rate. For example, resistance mutations may lead to a growth defect that manifests most strongly at low nutrient concentrations. Interestingly, the resistant phenotype (with higher maximal growth rate and lower enzyme affinity) is now playing the role of the debtor in a bacteriostatic environment, because it “grows now, pays later”. The debtor in the bacteriostatic environment mutates its enzymes to avoid the bacteriostatic effect, thereby sacrificing its future growth at low nutrient concentration because the mutated enzymes have lower affinity.

In our framework, growth penalties that are most severe in low nutrient conditions would manifest in the Monod affinity, *K*, increasing to *K* + Δ_*K*_, with a possible biophysical mechanism being mutations in proteins that reduce antibiotic binding but also reduce binding affinity for the natural substrate. (*28, 35*). Therefore, we shift our perspective and study the debt trade-off in which the debtor species has mutated so that it pays a growth cost in low-nutrient environment, but gains a maximal growth rate Δ*E*. How does this trade-off impact the value of debt?

We deduce the affinity-growth trade-off using an invasibility calculation: when would a minuscule amount of an ‘invader’ species succeed in expanding under nutrient dynamics dictated by a single dominant non-debtor? Namely, a species with lower enzyme affinity, *K* +Δ_*K*_, but also higher maximal growth rate, *E* + Δ_*E*_, is introduced to a system dominated by the single-species dynamics of an unperturbed (*K, E*) species. How large does Δ_*E*_ need to be to offset the cost of the decreased affinity *K*+Δ_*K*_? Equivalent growth implies that 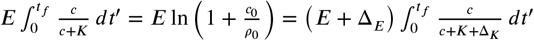 Expanding the perturbed Monod function at small Δ_*K*_ /*K*, with 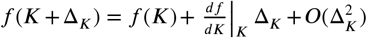, we write to leading order,

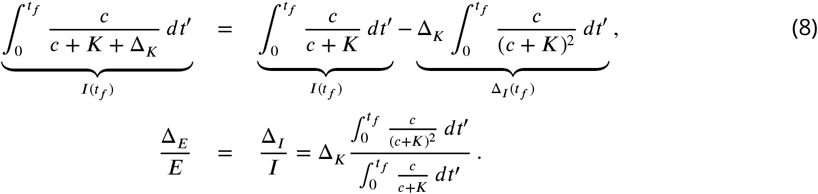

At equilibrium, the relative gain in maximal growth rate, Δ_*E*_/*E*, is offset by the relative loss in utilization, Δ_*I*_ /*I*. It is possible to approximate this relation at the low and high *c*_0_ limits, (see *SI Appendix* ***section 5***). The integral Δ_*I*_ and its approximate forms are shown in ***Figure 4***A. To leading order, we have,

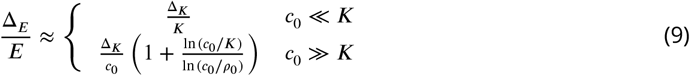

The relative cost at equilibrium behaves differently in the low *c*_0_ (chemostat) and high *c*_0_ limits. At low *c*_0_, as in a chemostat, the cost scales with Δ_*K*_. Therefore, for a given value of relative gain, Δ_*E*_/*E* there is a fixed cost Δ_*K*_ /*K* to maintain equilibrium, no matter how much nutrient is provided. There is little enough nutrient provided such that the growth function is linear in the nutrient amount, leaving the difference in affinity, Δ_*K*_, as the dominant term. A different picture emerges in the high *c*_0_ limit, where nutrient is initially saturating. Here, for a fixed gain, Δ_*E*_/*E* = const, the cost Δ_*K*_ ∼ *c*_0_, meaning that to maintain equilibrium, if more nutrient is provided, a larger enzyme affinity penalty can be supported. Said differently, the debtor takes over for a fixed Δ_*K*_ if *c*_0_ increases. The debtor advantage over the chemostat limit is shown in the horizontal tendency of the Δ_*K*_ /*c*_0_ boundary in ***Figure 4***B. This is because ‘payment’ happens mostly when *c* ≈ *K* + Δ_*K*_ and when *c*_0_ ≫ *K* and Δ_*K*_, a larger *c*_0_ gives the debtor an increasing span of growth with effectively no payment.

**Figure 4.**
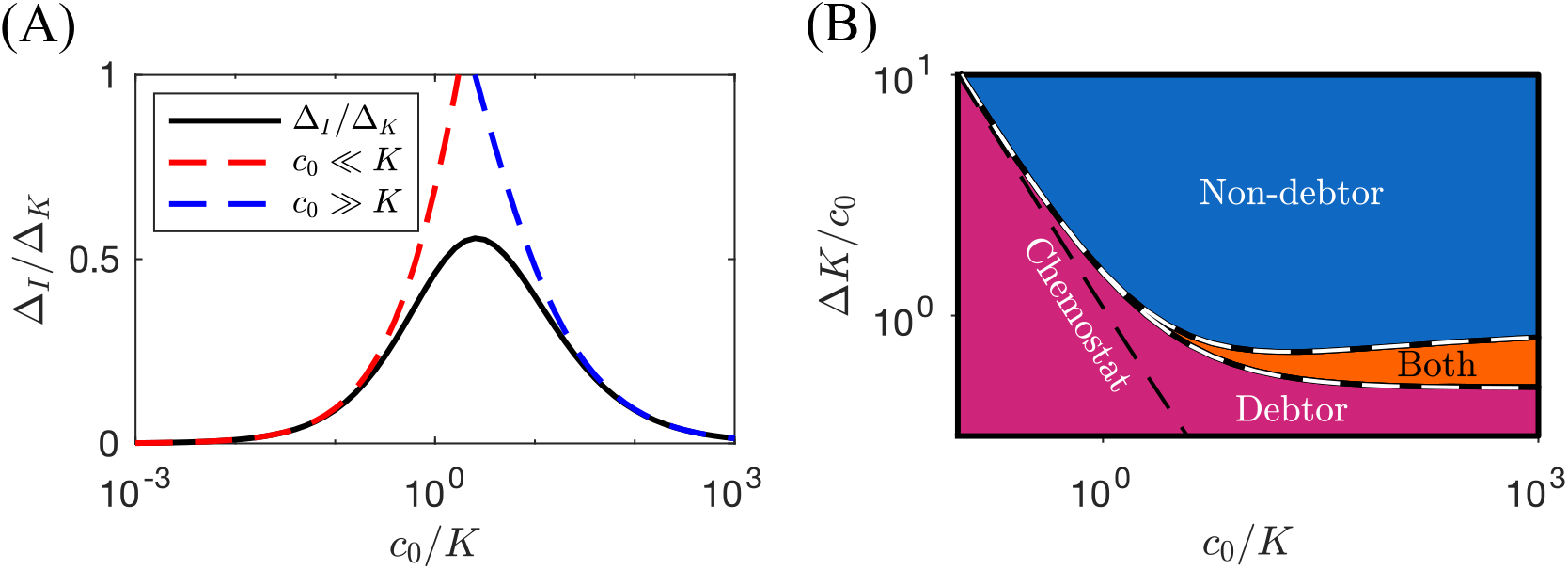
The tradeoff between maximal growth rate, *E* + Δ_*E*_ and reduced affinity due to *K* + Δ_*K*_. (*A*) Δ_*I*_ /Δ_*K*_ depicts the loss of growth per biomass, and its low and high *c*_0_/*K* approximations. (*B*) Phase diagram of the coexistence and dominance regimes, and its leading order analytic approximation (Eq. 9, white dashes). Black dashes: the chemostat limit of the phase boundary. *E* = Δ_*E*_ = 1 and *K* = 1. A coexistence region where both species support a nonzero population emerges high *c*_0_.

To complete the invasibility test, one must consider the opposite situation, where the debtor dominates the culture, dictating the nutrient dynamics with (*K* + Δ_*K*_, *E* + Δ_*E*_), and an invasion attempt by the non-debtor. When can the non-debtor invade? A similar derivation gives 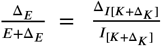, with the *I* and Δ_*I*_ integrals evolved under the debtor monoculture. As with the previous boundary, we may approximate, (*SI Appendix* ***section 5***). Interestingly, a coexistence region emerges at high *c*_0_/*K* (***Figure 4***B). Coexistence is due to the different effect of the dominant species: Δ_*E*_ is measured against *E* when the non-debtor is dominant but against *E* +Δ_*E*_ when the debtor is dominant. This mode of coexistence is similar to that seen in another recent serial dilution theory (*36*) and falls into a broader class of “relative nonlinearity” coexistence mechanisms (*37*).

### Stochastic stressor dynamics significantly lengthen lifetime of ecologically unstable species

The models explored in the prior sections exhibit temporal variability, but in a deterministic manner: dilution factors and stressor levels staying the same in all batch cycles. However, in real ecosystems the magnitude of the stressor effect is variable. For example, antibiotic concentration fluctuates in the environment (*15*), leaving a debtor species disproportionately susceptible to extreme dilution. What is the effect of such stochastic noise on the debt cost-benefit trade-off? Let Ω fluctuate, drawing Ω in each batch from a uniform distribution, Ω ∈ [Ω − Δ_Ω_, Ω + Δ_Ω_]. This corresponds to ‘pink’ or 1/Ω noise where extreme events occur at very low probability.

We consider a system with Ω > Ω^∗^ such that the debtor is outcompeted in the deterministic version of the model. We show this deterministic trajectory, batch to batch, plotting the relative abundance of the non-debtor (***Figure 5***A). To explore how this extinction process is affected by noise, we plot the stochastic trajectory averages of the equivalent system with two different levels of Ω noise. Intriguingly, the introduction of even a small level of noise, constituting 10% of Ω, significantly decelerates the extinction dynamics. With 10% noise, Ω ± 10%, the time to extinction is approximately tripled, while with 50% noise, the time to extinction is lengthened by orders of magnitude (***Figure 5***A). This behavior is symmetric to a reversal in roles: the same behavior occurs with non-debtor extinction if the environment favors debtors. Thus, noise in stressor severity can drastically extend the lifetime of ecologically unstable microbes.

**Figure 5.**
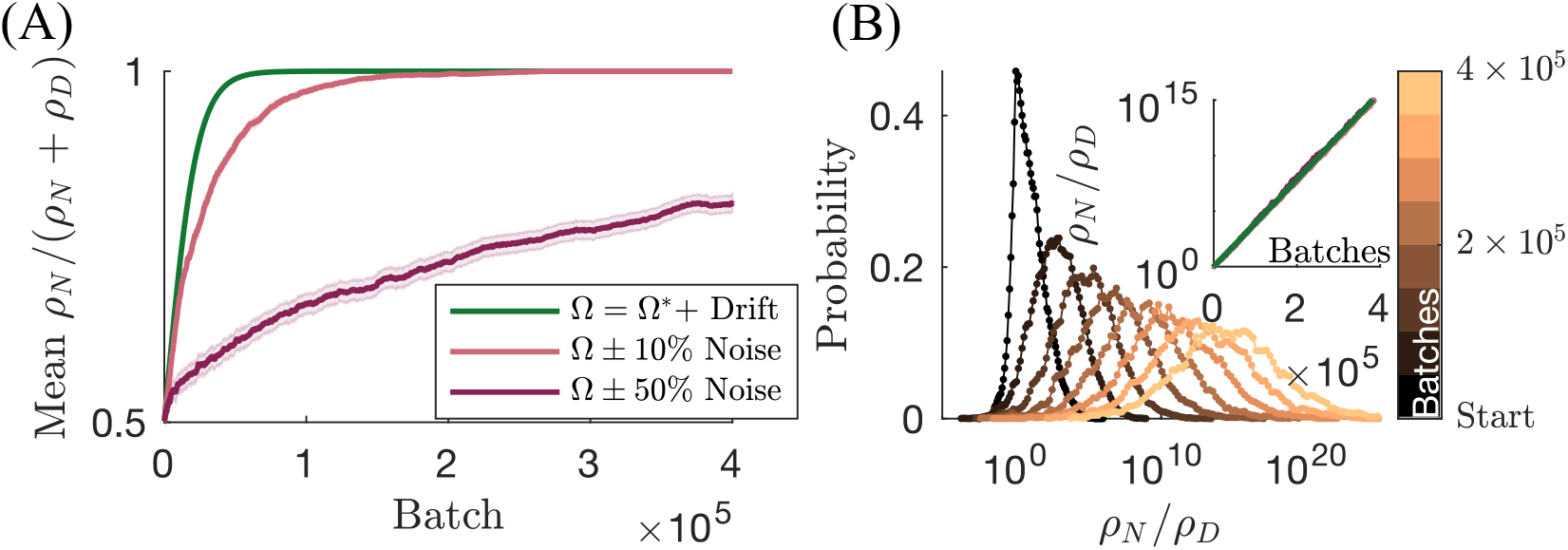
The effect of noise in Ω. (*A*) Full curves—mean *ρ*_*N*_ fraction. Shaded area: standard error of the mean, averaged over 1000 stochastic trajectories. Green curve—deterministic convergence to non-debtor take-over. (*B*) Time-series of the distribution of log_10_ *ρ*_*N*_ /*ρ*_*D*_, colors—distribution timeseries calculated over 50,000 batches, from the first batch until 4 × 10^5^ batches. The widening corresponds to a diffusion process and the drift the extinction of *ρ*_*D*_. Inset: the constant drift in the median value of *ρ*_*N*_ /*ρ*_*D*_, identical for all three noise levels, equivalent to the mode of the histograms.

How does this seemingly minor noise in Ω produce such drastic departures from the deterministic behavior? The key observation is that Ω *t*_*f*_ (and *ω t*_*f*_) act on the population abundances in logarithmic space (***Equation 3*** and ***Equation 5***). Thus, rather than inducing a random walk of *ρ*_*N*_ in linear space (which would preserve the deterministic dynamics of the mean), this process is instead a random walk in log(*ρ*_*N*_ /*ρ*_*D*_). In ***Figure 5***B we plot the evolution of the distribution of the *balance* variable, log(*ρ*_*N*_ /*ρ*_*D*_) (*38*). As can be seen, the balance undergoes a canonical biased random walk, with the distribution widening and simultaneously its mean drifting. The drift is due to the induced extinction because unfavorable debt conditions, Ω > Ω^∗^, were chosen. As expected, the mean log(*ρ*_*N*_ /*ρ*_*D*_) of the different noise levels, including the deterministic dynamics, are all equal (***Figure 5***B inset). As is known, the balance or log-ratio is the appropriate degree of freedom to consider when studying the dynamics of a composition (*39, 40*). Indeed, balances would be the appropriate variables to consider in multi-species generalization of our framework. This random walk in logarithmic space has long tails for the relative abundances, 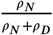, a quantity typically relevant ecologically and measured in experiments. Thus, large, rare events significantly influence an observed mean fraction even when noise is relatively small. This suggests a broad distribution of stress resistance traits even within a single species, consistent with what has been observed in *E. coli* (*12*).

## Conclusions and Discussion

Real-world microbes can incur debt by increasing immediate growth while neglecting protection from future events. Here, we proposed a theory for the impact of such debt trade-offs on microbial competition. We began by formulating chemostat debt dynamics and validating them with data from a classical chemostat experiment, thereby exposing the dynamic debt regulation behavior of *E. coli*. Notably, we validated the predicted linear relationship between the inverse dilution rate and the debtor advantage proposed by our chemostat theory. Having established the basic debt dynamics, we extended our theory to serial-dilution cultures. We found that growth debt where the ‘payment’ occurs continuously and where it occurs at the end of the batch are equivalent and can be unified in a single framework. We then constructed a universal phase diagram that delineates the boundary where debt becomes competitively favorable, as a function of nutrient amount in the system. This phase diagram encompasses both serial dilution and chemostat dynamics, unifying the two experimental systems. In the chemostat, the trade-off line is proportional to the total nutrient supply. Conversely, in the serial dilution setup, the addition of nutrients in a saturating concentration results in a diminishing return on debt. The effect of diminished return can be considerable: a non-debtor that would typically go extinct in a high-nutrient chemostat may dominate an equivalent serial dilution culture. However, we found that not all forms of debt are the same: debt associated with enzyme affinity allows the coexistence of two species on a single resource, violating the competitive exclusion principle by forming temporal niches (*36*).

Experiments to test the predicted phase boundaries would involve competitions between a debtor and non-debtor strain in varying serial dilution environments. One could monitor the persistence of an antibiotic resistance element in a community across a range of antibiotic and nutrient concentrations (thus varying *ω* and *c*_0_/*K*) to construct an empirical version of the phase diagram in ***Figure 3***A. The debtor could be a wild-type *E. coli* with the non-debtor being that same strain with an antibiotic resistance element. Ideally this element would confer resistance that does not inactivate the antibiotic (e.g. an efflux transporter), to prevent benefit to non-resistant organisms (*41*). The resulting phase diagram should feature a boundary defined by ***Equation 7***. To construct a theoretical phase diagram for comparison, one could measure death kinetics as a function of antibiotic concentration, and in the absence of antibiotic use a competition experiment to infer the cost of the element. In the chemostat analysis (***Figure 1***), we demonstrated how to estimate both values given a set of experimental data. Evolution may complicate such an experiment—discussed below.

Stochastic stressor dynamics, a realistic extension of the model, revealed unexpected behavior. Even small fluctuations in stress level can significantly delay the extinction of a species due to the compositional nature of the serial-dilution process. To decouple the compositional effects, a log-ratio is used (*38–40*), here, log 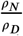. As the system relaxes to steady state, the mean log-ratio dynamics are same in the deterministic and stochastic systems. Conversely, the mean relative fraction 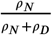 changes greatly with the addition of noise, leading to delayed extinctions. The outcomes of our stochastic dynamics study hold considerable implications for the management and control of antibiotic resistance elements within environments. For example, rotating antibiotics to reduce the prevalence of resistance elements has been proposed as a means of limiting resistance proliferation (*42*). Our findings indicate that the timescale for the depletion of resistance elements in the environment may be significantly longer than deterministic models (*43*) would estimate, limiting the efficacy of rotation-based strategies. Our findings also extend beyond antibiotic resistance, implying that stress resistance genes could persist within communities for extended periods despite a net loss of fitness. Noisy debt dynamics may broadly influence the prevalence of accessory genes in bacteria.

The predictions on the influence of stressor noise on the lifespan of resistance elements can be experimentally tested. The model predicts that across many independent populations of cells, populations subject to a more noisy stressor are expected to have longer mean persistence of resistance elements (***Figure 5***). One approach is to first identify a constant stressor level at which a resistance element goes extinct over multiple serial passages. For an antibiotic resistance element, this would be a sub-MIC level of antibiotic that impedes growth, but does not impose a large enough growth defect to allow persistence of the element. With baseline stressor environment established, one can run high-throughput batch cultures, each receiving a randomized amount of antibiotic each passage, and with control wells receiving the same mean amount). Resistance could be measured as a population mean, as in Notley-McRobb *et al*. (*11*). The model predicts that the mean level of resistance in the noisy communities will exceed that in the constant-stressor controls. Even in the presence of additional mechanisms evolving over time, the impact of noise on overall resistance levels should still be observable through community-wide assays. The distribution of stress resistance levels observed in this experiment could also be compared to those found in nature (*12*).

Our theory has explored only one form of stressor fluctuations. In addition to inter-batch stress level changes, stressor intensity may also change during the batch, e.g., antibiotics can be produced by other microbes, resulting in nontrivial stressor dynamics (for a chemostat analysis of these dynamics, see (*25*)). This may lead to large departures from our predicted behaviors: for example, such anti-symmetric microbial interactions have been found in theory to lead to rock-paper-scissor dynamics (*44*). Furthermore, we have not considered the case where the dead bacteria contribute nutrient to the environment. Allowing such recycling may weaken the debtor advantage since instead of depriving the competitor of nutrient the debtor would then simply store and then release the nutrient. In summary, while this manuscript covers the impact of exogenous stressors, to elucidate the impact of endogenous stressors on debt will require further work.

Focusing on the fundamental mechanisms underlying microbial debt, we considered only two species and a single nutrient. Yet, natural microbial communities typically feature many coexisting species. The mortality (*ω*, Ω) debt trade-off is not capable of supporting two species, so it would not support multiple species coexistence. Conversely, the affinity debt trade-off may support more than two species coexisting but may require fine-tuning (*36*). Investigating debt dynamics within complex, randomly assembled ecosystems (*45*) may shed light on the functioning of multispecies communities, such as the gut microbiome. Natural microbial communities also tend to feature more than the single resource assumed in our model. Previously, we demonstrated the ‘early-bird’ effect, where the presence of multiple resources can amplify the benefit of an early growth advantage within a batch (*20*). This ‘early-bird’ microbe can leverage its larger population size later in the batch to deplete resources it may not specialize in, depriving competitors. Thus, we expect debt dynamics to influence ecosystems with multiple species and resources in nontrivial ways.

Evolution is an important factor to consider, as the emergence of new variants of debtor and non-debtor occurs by mutation and subsequent competition with their ancestors. A promising direction to analyze the evolution of debt strategies is by using adaptive dynamics (*46*). Evolution may lead to the dominance of a single strain in the chemostat context, as it tends to minimize the steady-state nutrient level— the phenomenon of ‘pessimization’ observed widely in adaptive dynamics (*46*). Pessimization is consistent with the pH 7 chemostat results of Notley-McRobb *et al*. (*11*), where the debtor dominates. At other pH values, partial *rpoS* mutants emerged and the chemostat did not reach steady state, so we cannot conclude whether the pessimization principle held in these cases. Note that later work from the same authors demonstrated that it is possible to qualitatively predict the outcomes of such evolution experiments using a chemostat theory (*27*). In a serial dilution context, it unclear whether the chemostat ‘pessimization’ intuition holds, particularly as the switch from chemostat to serial dilution also can introduces additional stressors. For example, microbes can substantially acidify growth media, exposing the community to stress at the end of the batch (*47*). Such pH stress effectively increases the cost of debt beyond what is currently in our model. Additionally, close matching of these evolution data may require additional information on the shape of the debt trade-off surface (*9, 27*), outside the scope of this manuscript.

Through the ages, debt has been an ever-present part of human life (*48*). Yet the omnipresence of debt in human life mirrors a much older and more basic notion. When competition for resources is fierce and resource levels change with time, a “grow now, pay later” strategy can be exploited to gain an advantage over one’s competitors, a manifestation of the “early-bird” effect (*20*). Such an early-bird strategy is built on depriving competitors of resources by growing quickly, even if it means paying a significant future cost. Our work has shed light on how such fundamental tradeoffs influence the world.

## Acknowledgments

This work was partially funded by AE’s startup grant at the Hebrew University. Both JGL and AE conceived the idea, carrier out calculations, and wrote the manuscript. We thank Nathalie Balaban, Po-Yi Ho, and Anthony Lyndon Shiver for critically reading this manuscript.

The authors declare that they have no competing interests.

All code and data are available at github.com/AmirErez/MicrobialDebt.

## Appendix 1

### Chemostat debt trade-offs

We consider a single debtor species growing within a chemostat and compute the values of Δ_*E*_ and *ω* at which this species has equivalent competitive ability as a non-debtor species.

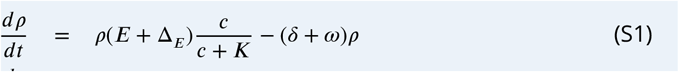

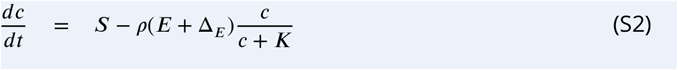

The outcome of competition within a chemostat is determined by the steady-state nutrient concentration, *c*^∗^. Species that can survive at the lower *c*^∗^ will invade and exclude species with higher *c*^∗^. From Eq. S1, the *c*^∗^ for a debtor species will be

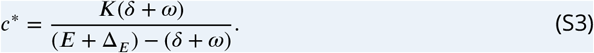

We now determine the values for Δ_*E*_ and *ω* at which the debtor has the same *c*^∗^ as a non-debtor:

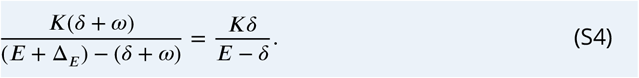

Solving the above expression yields:

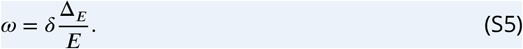

Thus, in a chemostat, a growth benefit of Δ_*E*_ is worthwhile if it comes with 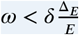

Note that the requirement *δ* < *E* must be satisfied otherwise the chemostat washes out. Therefore, if *c*_0_ increases, the steady state *ρ* must also increase, since increasing the nutrient concentration S while keeping the dilution rate constant gives *S*/*δ* = *ρ*.

## Appendix 2

### Chemostat model fitting to experimental data

To demonstrate applying our theory to empirical population dynamics, we fit data from Fig. 1A of Notley-McRobb et al. (*11*). We focus on these data as the chemostats at other pH values did not reach steady state. We first digitally extracted data from the figures. We utilize the chemostat equations in Eq. S2 to model these data. As these populations are not exposed to external stress, we set *ω* = 0. Chemostats near steady state operate in a low-nutrient regime, so we approximate the growth function to be linear 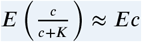, absorbing the factor 1/*K* into *E*. Chemostats approach nutrient steady-state quickly, and thus for simplicity we make a separation of timescales approximation, assuming the nutrients are pseudo-steady-state relative to the bacterial dynamics. Since the dominant source of nutrient depletion is microbial consumption, we neglect nutrient losses due to dilution. We do not explicitly model mutation, but rather assume mutants arise very early in the system and fit an initial relative abundance of the non-debtor species *x*_*D*_(*t* = 0). Dilution rate *δ* is set based on the specified chemostat dilution rate. To convert from “generations” to physical time, we multiply by ln(2)/*δ*. Due to the fact that the input data are relative abundances, we can only estimate Δ*E*/*E* and also set *S* = 1. Δ*E*/*E* was estimated separately for each dilution. The model fitting and confidence/prediction interval estimation was performed using the same methods as in (*20*).

To apply our data to stressor deal kinetics, we fit data from Fig. 3A of Notley-McRobb et al., digitally extracting the data as above. We model the surviving fraction of the total microbial population as *f* = (1 − *x*_*N*_ (*t* = 0)) exp(−*ω*_*D*_*t*) + *x*_*N*_ (*t* = 0) exp(−*ω*_*N*_ *t*), where *x*_*N*_ (*t* = 0) is set using measurements from Notley-McRobb et al. All data is fit using a single value of *ω*_*N*_ and *ω*_*D*_. Fitting and confidence/prediction interval estimation is performed as above. Note, however, that our current theory is not able to model all stressors: Notley-McRobb *et al*. also subjected their cultures to hydrogen peroxide stress and the resulting cell death kinetics deviate substantially from the first-order kinetics we assume.

## Appendix 3

### Serial dilution model dynamics in dimensionless variables

It is sometimes useful to cast the dynamics in dimensionless variables. Our starting point is:

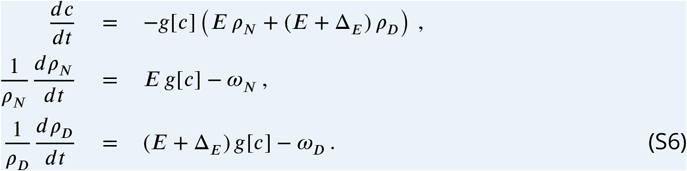

By defining time in units of the growth rate of the normal non-debtor, 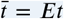, similarly, nutrient in units of 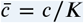. We annotate 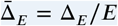 and 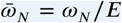. Explicitly using the Monod form for *g*[*c*] we have,

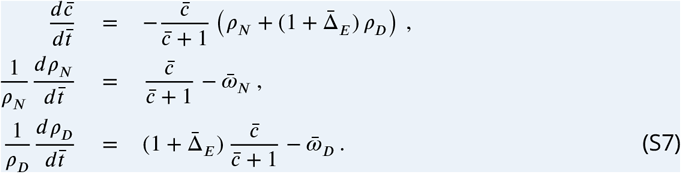

## Appendix 4

### Derivation of the chemostat limit of the serial dilutions, *c*_0_ ≪ *K*

Let species N be the normal, non-debtor species, growing at maximal rate *E*, and diluted by 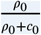 to the next batch. Let the debtor species D grow at maximal rate *E* + Δ_*E*_ and get diluted by 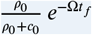. We expand the low-nutrient limit,

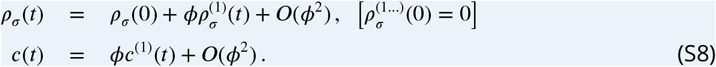

We note that in the *c*_0_/*K* ≪ 1 limit (i.e., *ϕ* ≪ 1), we have,

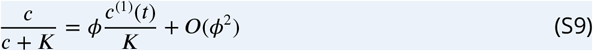

The nutrient dynamics to order *ϕ* are,

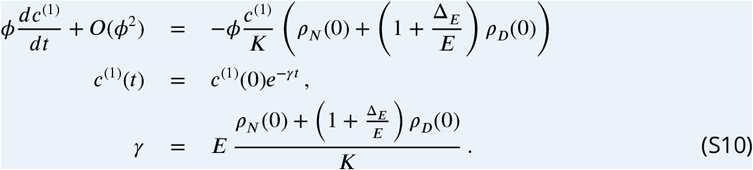

We may now solve for the time-course integral,

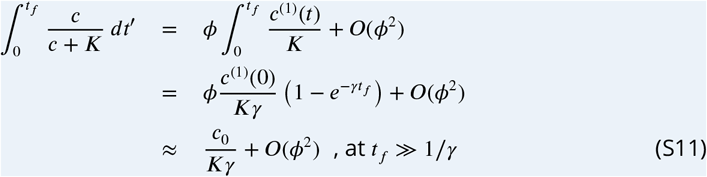

The species abundance dynamics follow,

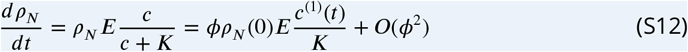

Matching order *ϕ* and integrating both sides until time *t*_*f*_, and using 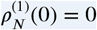, gives us

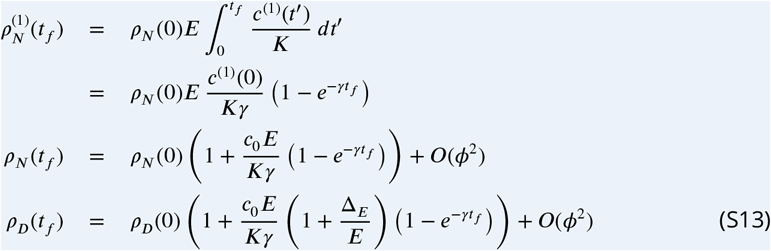

At time (*t*_*f*_) in batch *b*, each species gets diluted by a different fraction. Therefore, at the beginning of batch *b* + 1,

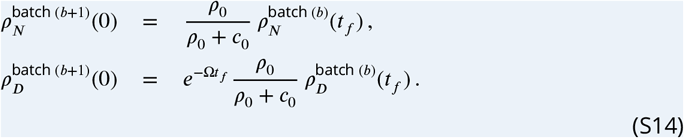

We require steady state, meaning that 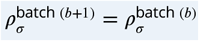.This gives,

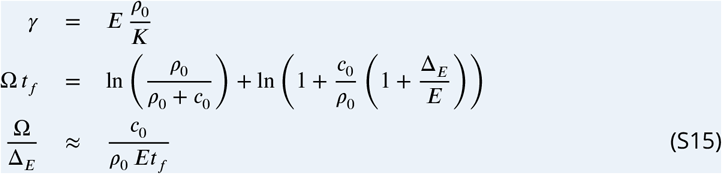

Therefore, Ω/Δ_*E*_ ∼ *c*_0_ in the chemostat limit, *c*_0_ ≪ *K, ρ*_0_ as shown in ***Figure 3***.

In the language of the chemostat, the steady state requires 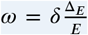 (***Equation S5***). In the serial dilution framework, 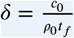 and 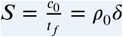 (*20*), therefore,

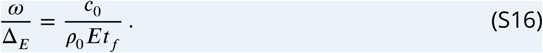

This is precisely the relation derived in the leading-order *c*_0_/*K* expansion, Eq. S15.

## Appendix 5

### Debt trade-offs with differing dilution in the limit of high *c*_0_

Here, we compute the growth-debt trade-offs in the limit of high *c*_0_. In this scenario, the *c*(*t*) ≫ *K* during the vast majority of growth. Thus, we approximate 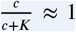 in this computation, a unit step function that suddenly terminates and gives zero. This approximation yields the following set of ODEs governing the batch dynamics:

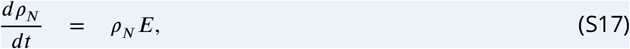

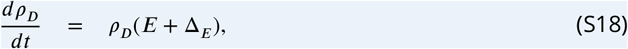

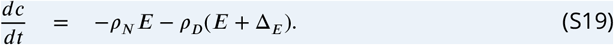

Both populations are therefore described by simple exponential functions, *ρ*_*N*_ (*t*) = *ρ*_*N*_ (0)*e*^*Et*^ and 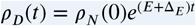. Substituting these functions into the nutrient equation and solving yields 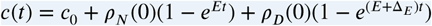.

### Response to Δ_*K*_

We may evolve *c*(*t*) according to the single species dynamics and calculate the definite integral. If the single species is a non-debtor, at high *c*_0_/*K*, we have 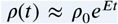 and therefore, *c*(*t*) = *c*_0_ + *ρ*_0_(1 − *e*^*Et*^). Accordingly, 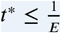 ln 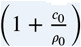.

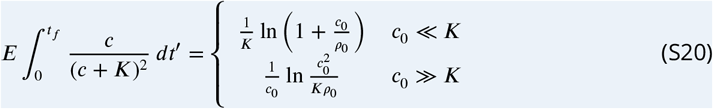

Therefore, at low *c*_0_/*K* we have ln(1 + *c*_0_/*ρ*_0_) ≈ *c*_0_/*ρ*_0_ and,

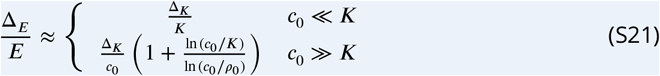

Similarly, we may consider the opposite situation, where we evolve under the debtor species, (*K* + Δ_*K*_, *E* + Δ_*E*_), and find the condition that the normal species can invade. A similar derivation gives 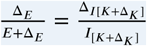, with the *I* and Δ_*I*_ integrals evolved under the debtor species. Again, we may approximate, 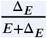ln 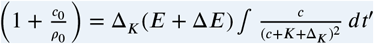, giving,

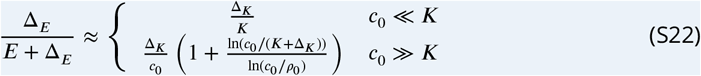

#### The chemostat equivalent for a *K* debtor

We can consider the chemostat equivalent directly. We consider a single debtor species growing within a chemostat and compute the values of Δ_*E*_ and Δ_*K*_ at which this species has equivalent competitive ability as a non-debtor species.

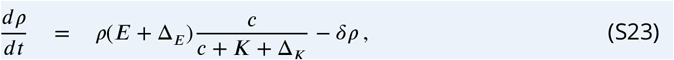

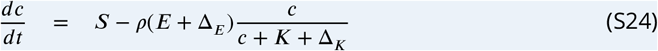

The equilibrium for a debtor with the *c*^∗^ of a non-debtor is,

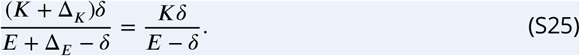

Solving the above expression yields 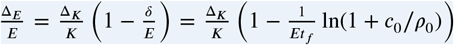. In the language of the serial-dilutions framework, 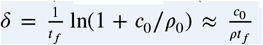 and 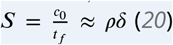, therefore,

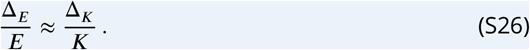

This is the relation derived in the leading-order *c*_0_/*K* expansion.

**Appendix 6—figure S1.**
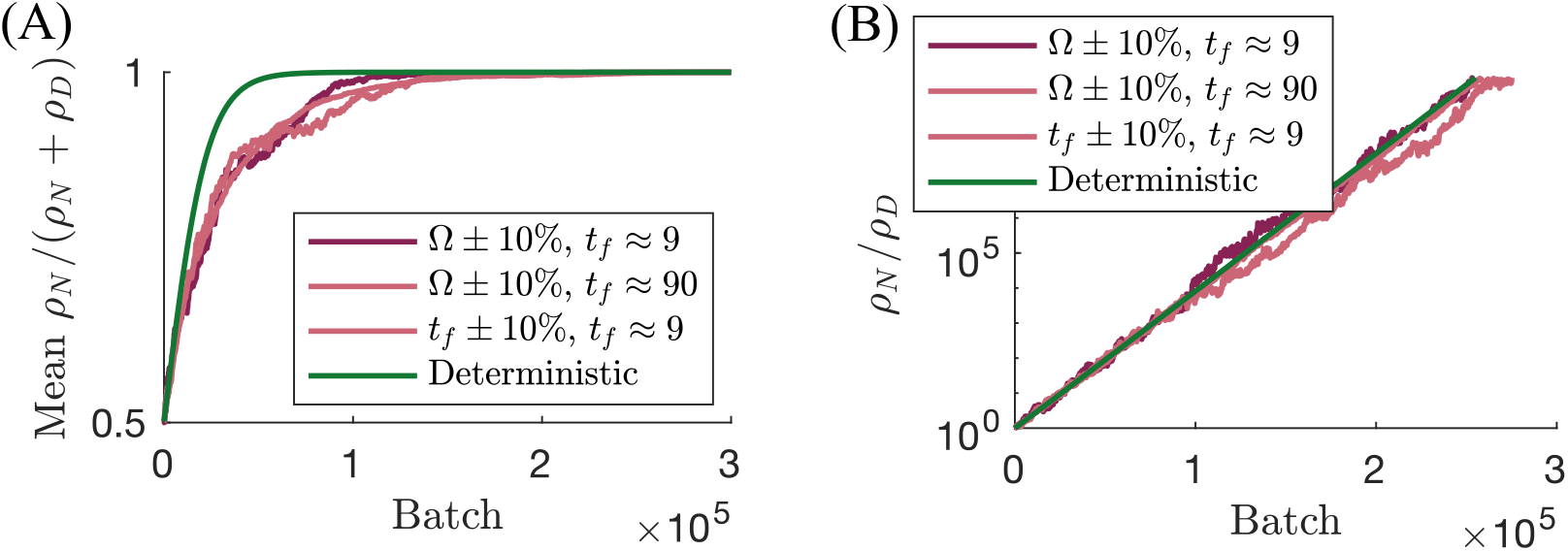
Comparing the effect of noise in *t*_*f*_ and Ω. (*a*) Full curves - mean *ρ*_*N*_ fraction, averaged over 1000 stochastic trajectories. Green curve - deterministic convergence to non-debtor take-over. The three stochastic regimes show a similar, and much slower trend to mean *ρ*_*N*_ taking over the population than the deterministic case. (*b*) The median relative species log-fold-difference *ρ*_*N*_ /*ρ*_*D*_. Viewed this way, the drift towards *ρ*_*N*_ takeover is the same for both the deterministic and the three stochastic regimes.

## Appendix 6

### Comparing noise in *t*_*f*_ with noise in Ω

If we keep all conditions the same, fluctuations in *t*_*f*_ are different from fluctuations in Ω because *t*_*f*_ affects both species through the utilization integral. We sought out to understand if fluctuations in *t*_*f*_ therefore lead to different behavior. We simulated a system that drifts to the debtor going extinct, by imposing a small positive Ω on top of the equilibrium Ω value that fits the debtor’s 5% gain in maximal growth rate. ***Figure S1*** compares the effect of noise in *t*_*f*_ with noise in Ω. As is apparent, the noise in *t*_*f*_ acts the same way.

## References

1. J. Lloyd-Price, G. Abu-Ali, C. Huttenhower, Genome Medicine 8, 51 (2016).

2. A. Crits-Christoph et al., Microbiome 1, 28 (2013).

3. S. Lladó, R. López-Mondéjar, P. Baldrian, Microbiology and Molecular Biology Reviews 81, e00063–16 (2017).

4. L. W. Kelly, A. F. Haas, C. E. Nelson, mSystems 3, e00162–17, eprint: 10.1128/mSystems.00162-17 (2018).

5. F. Abram, T. Arcari, D. Guerreiro, C. P. O’Byrne, in ed. by R. K. Poole, D. J. Kelly (Academic Press, 2021), vol. 79, pp. 133–162.

6. M. Linkevicius, J. M. Anderssen, L. Sandegren, D. I. Andersson, Journal of Antimicrobial Chemotherapy 71, 1307–1313 (2016).

7. V. Patel, N. Matange, eLife 10, e70931 (2021).

8. A. H. Melnyk, A. Wong, R. Kassen, Evolutionary applications 8, 273–283 (2015).

9. K. Phan, T. Ferenci, The ISME journal 11, 1472–1482 (2017).

10. S. Nair et al., Antimicrobial Agents and Chemotherapy 66, e01529–21 (2022).

11. L. Notley-McRobb, T. King, T. Ferenci, Journal of bacteriology 184, 806–811 (2002).

12. E. Y. Valencia, J. P. Barros, T. Ferenci, B. Spira, Microbial ecology 83, 68–82 (2022).

13. S. A. Smits et al., Science 357, 802–806 (2017).

14. J. F. Brooks et al., Cell 184, 4154–4167.e12 (2021).

15. V. Diwan, C. S. Lundborg, A. J. Tamhankar, PLOS ONE 8, e68715 (2013).

16. I. Cvijović, B. H. Good, E. R. Jerison, M. M. Desai, Proceedings of the National Academy of Sciences 112, E5021–E5028 (2015).

17. J. Monod, Recherches sur la croissance des cultures bacteériennes (Hermann and Cie, 1942).

18. J. E. Goldford et al., Science 361, 469 (2018).

19. N. Meroz, N. Tovi, Y. Sorokin, J. Friedman, Nature Communications 12, 2891 (2021).

20. A. Erez, J. G. Lopez, B. G. Weiner, Y. Meir, N. S. Wingreen, eLife 9, ed. by D. Weigel, A. Sanchez, A. Succurro, L. Dai, e57790 (2020).

21. A. Erez, J. G. Lopez, Y. Meir, N. S. Wingreen, Physical Review E 104, 044412 (2021).

22. P.-Y. Ho, B. H. Good, K. C. Huang, eLife 11, ed. by N. Segata, W. S. Garrett, S. Gibbons, e75168 (2022).

23. L. Pacciani-Mori, A. Giometto, S. Suweis, A. Maritan, PLOS Computational Biology 16, e1007896 (2020).

24. A. Aranda-Diaz et al., bioRxiv, 2023–01 (2023).

25. S.-B. Hsu, P. Waltman, Mathematical biosciences 187, 53–91 (2004).

26. R. E. Lenski, S. E. Hattingh, Journal of Theoretical Biology 122, 83–93 (1986).

27. R. Maharjan et al., Ecology letters 16, 1267–1276 (2013).

28. A. D. Letten, A. R. Hall, J. M. Levine, Nature ecology & evolution 5, 431–441 (2021).

29. M. Basan et al., Nature 584, 470–474 (2020).

30. E. Kussell, S. Leibler, Science 309, 2075–2078 (2005).

31. J. Lin, M. Manhart, A. Amir, Genetics 215, 767–777 (2020).

32. K. Phan, T. Ferenci, The ISME Journal 7, 2034–2043 (2013).

33. Y. Kaplan et al., Nature 600, 290–294 (2021).

34. A. Posfai, T. Taillefumier, N. S. Wingreen, Phys. Rev. Lett. 118, 028103 (2 2017).

35. T. Song et al., Molecular microbiology 91, 1106–1119 (2014).

36. Y. Fridman, Z. Wang, S. Maslov, A. Goyal, PLOS Computational Biology 18, e1010244 (2022).

37. P. Chesson, Theoretical population biology 45, 227–276 (1994).

38. J. Rivera-Pinto et al., mSystems 3, e00053–18 (2018).

39. J. Aitchison, Journal of the Royal Statistical Society: Series B (Methodological) 44, 139–160 (1982).

40. J. Friedman, E. J. Alm, PLOS Computational Biology 8, e1002687 (2012).

41. R. A. Sorg et al., PLoS biology 14, e2000631 (2016).

42. E. M. Brown, D. Nathwani, Journal of antimicrobial chemotherapy 55, 6–9 (2005).

43. G. F. Webb, E. M. D’Agata, P. Magal, S. Ruan, Proceedings of the National Academy of Sciences 102, 13343–13348 (2005).

44. E. D. Kelsic, J. Zhao, K. Vetsigian, R. Kishony, Nature 521, 516–519 (2015).

45. R. Marsland III et al., PLoS computational biology 15, e1006793 (2019).

46. O. Diekmann, BANACH CENTER PUBLICATIONS 63, 47–86 (2002).

47. C. Ratzke, J. Denk, J. Gore, Nature ecology & evolution 2, 867–872 (2018).

48. D. Graeber, Debt: The First 5,000 Years (Melville House, BROOKLYN, NY, Reprint edition, 2012).

